# Deletion of LRP1 in Smooth Muscle Cells Induces Infrarenal Aortic Aneurysms in Mice

**DOI:** 10.64898/2026.10.16.759442

**Authors:** Michael K. Franklin, Guan Fan, Bowen Li, Pengjun Wang, Deborah A. Howatt, Jessica J. Moorleghen, Adam E. Mullick, Alan Daugherty, Hong S. Lu

**Affiliations:** Saha Cardiovascular Research Center and Saha Aortic Center, University of Kentucky, Lexington, Kentucky, USA; Department of Physiology, University of Kentucky, Lexington, Kentucky, USA; Ionis Pharmaceuticals, Inc., Carlsbad, CA, USA

**Keywords:** Low-density lipoprotein receptor-related protein 1, smooth muscle cells, aorta, aortic aneurysm, angiotensinogen, antisense oligonucleotides, micro-CT

## Abstract

**Background:** Low-density lipoprotein receptor-related protein 1 (LRP1) in smooth muscle cells (SMCs) is essential for maintaining aortic wall integrity. This study investigated whether deletion of LRP1 in SMCs induces region-specific aortic pathology and evaluated a potential therapeutic strategy.

**Methods:** *Lrp1*-floxed mice were bred with *Sm22-Cre*+/0 mice to generate mice with SMC-LRP1 deletion (SMC-LRP1-/-) and wild-type littermates (SMC-LRP1+/+). Micro-computed tomography (micro-CT) was used to visualize the aorta, and aortic tissue sections were characterized histologically. To evaluate a potential therapeutic strategy, antisense oligonucleotides (ASOs) targeting hepatocyte-derived angiotensinogen (AGT) were administered to inhibit AGT synthesis.

**Results:** Immunostaining and Western blotting confirmed LRP1 deletion in aortic SMCs. At 40 weeks of age, micro-CT revealed marked increases in aortic width and volume. Although dilatation occurred throughout the aorta, the infrarenal region exhibited the greatest expansion in both male and female SMC-LRP1-/-mice compared with SMC-LRP1+/+ littermates; subsequent analyses therefore focused on this region. LRP1 deletion in SMCs caused pronounced SMC loss and increased collagen deposition in the infrarenal aorta. Inhibition of hepatocyte-derived AGT with ASO attenuated infrarenal aortic dilatation and preserved aortic wall integrity in both male and female mice.

**Conclusions:** Deletion of LRP1 in SMCs causes progressive aortic dilatation that includes the infrarenal aorta. Pharmacologic inhibition of hepatocyte-derived AGT attenuates infrarenal aortic dilatation.

**What Are the Clinical Implications?:** Deletion of low-density lipoprotein receptor-related protein 1 (LRP1) in smooth muscle cells (SMCs) causes diffuse aortic dilatation, with spontaneous aneurysm formation most prominent in the infrarenal abdominal aorta—the most common site of abdominal aortic aneurysms (AAAs) in humans. Given the strong association between LRP1 variants and human AAAs, these findings identify SMC-derived LRP1 as a critical contributor to aortic wall integrity and provide a valuable tool for investigating disease mechanisms. Importantly, pharmacological inhibition of hepatocyte-derived angiotensinogen with an antisense oligonucleotide significantly attenuated infrarenal aortic dilatation, highlighting a promising therapeutic strategy for AAAs.

**GRAPHIC ABSTRACT:** 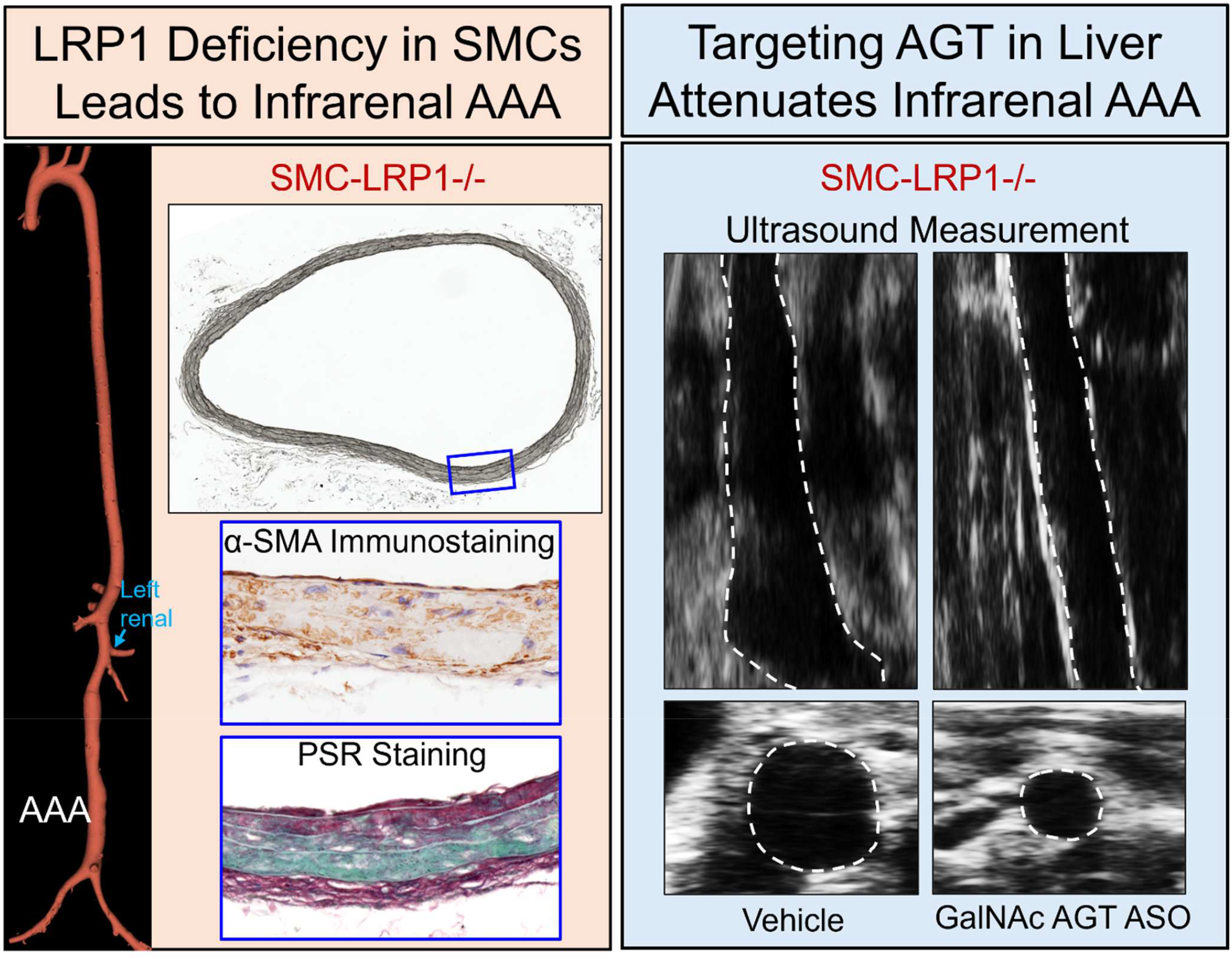

## INTRODUCTION

Low-density lipoprotein receptor-related protein 1 (LRP1) is a multifunctional protein that binds a broad array of ligands, many of which are proteases.^1^A major function of LRP1 is the endocytosis of protease ligands. In the vasculature, the removal of extracellular proteases by smooth muscle cells (SMCs) is presumed to be critical for maintaining the structural integrity of the vascular wall.^1, 2^

SMCs are the most abundant cell type in the aortic wall, and LRP1 is enriched in SMCs.^3^Several studies has demonstrated that genetic deletion of *Lrp1* in SMCs leads to aortic pathologies in mice.^4-8^Consistent with these findings in mice, genome-wide association studies (GWAS) have identified variants in the *LRP1* gene associated with abdominal aortic aneurysms (AAAs).^9-11^The role of LRP1 in human AAAs is further supported by the reduced abundance of LRP1 protein in AAA tissue.^12^Moreover, genetic variants in *LRP1* have been linked to acute thoracic aortic dissection,^13^reinforcing its critical role in preserving aortic wall integrity and preventing catastrophic aortic complications. Collectively, these findings underscore the importance of LRP1 in maintaining the structural and functional integrity throughout the aortic wall.

Previous studies in mice with deletion of LRP1 in SMCs (SMC-LRP1-/-) have reported spontaneous, progressive dilatations of the aortic root^4^and the superior mesenteric arteries^8^by 30-40 weeks of age. Given the previously reported pathological changes in the aortic root and ascending aorta of SMC-LRP1-/-mice, we first performed micro-computed tomography (micro-CT) imaging to visualize the entire aorta in mice at 40 weeks of age, which corresponds approximately to 40 years of age in humans. As expected, we observed thoracic aortic dilatations. Unexpectedly, we also detected dilatation of the infrarenal aorta, the region most commonly affected by AAAs in humans, but not typically involved in angiotensin II (AngII) infusion-induced AAAs in mice.^14, 15^To investigate this unique AAA phenotype in mice with SMC-LRP1 deletion, we performed histological characterization and determined whether inhibition of angiotensinogen (AGT), the sole precursor of AngII, attenuated infrarenal aortic dilatation.

## METHODS

### Data Availability

Detailed material information is available in the Major Resources Tables. Numerical data are available in the Supplemental Data File. Raw data and related analysis supporting the findings of this study are also available from the corresponding authors upon reasonable request.

### Mice

Female *Lrp1* f/f mice (Strain #: 012604; The Jackson Laboratory) were bred with male *Sm22*-*Cre*+/0 mice (Strain #: 017491; The Jackson Laboratory). The breeding strategy for generating *Sm22-Cre*0/0*;Lrp1* f/f (SMC-LRP1+/+) and *Sm22-*Cre+/0*;Lrp1* f/f (SMC-LRP1-/-) is shown in Figure S1A and Major Resources Tables.^5, 8^*Cre* genotyping was performed by polymerase chain reaction on tail-tip DNA. Reduction of the LRP1 protein in the aortic media was confirmed by both Western blotting and immunostaining.

All mice were maintained in a barrier facility on a light:dark cycle of 14:10 hours and fed a normal laboratory diet (Diet #: 2918; Inotiv) after weaning. Both male and female littermates were used for the experiments reported in this manuscript in accordance with the Statement from the ATVB Council.^16^At termination, mice were euthanized using an overdose of a ketamine and xylazine (90 and 10 mg/kg, respectively) cocktail. All animal experiments were performed according to protocols approved by the University of Kentucky Institutional Animal Care and Use Committee (Protocol number 2018-2967).

### Antisense Oligonucleotides Targeting Hepatocyte-derived Angiotensinogen

Mouse GalNAc AGT ASO (5’-ATCATTTATTCTCGGT-3’)^17^was provided by Ionis Pharmaceuticals Inc. (Carlsbad, CA, USA). Four-week-old male and female SMC-LRP1-/-mice were injected subcutaneously with PBS (vehicle) or GalNAc AGT ASO (5 mg/kg) once weekly until 40 weeks of age. Vehicle-injected SMC-LRP1+/+ mice were used as normal controls because they do not develop infrarenal aortic aneurysms. At termination, plasma was collected to measure AGT concentrations using a mouse AGT ELISA kit (Catalog #: ab245718; Abcam).

### Micro-CT Scanning and 3D Reconstruction

Micro-CT was performed as described previously.^8, 18^Mice were euthanized with an overdose of ketamine and xylazine. The thoracic cavity was cut open, and the right atrium was incised to permit blood drainage. The vasculature was perfused with saline (5 mL) through the left ventricle.

Immediately thereafter, the right atrium was sealed with cyanoacrylate adhesive, and Microfil (Flow Tech, Inc.) or Vascupaint (MediLumine) was injected through the same catheter. Once the contrast agent became visible in the arterioles surrounding the small intestine, the catheter was clamped to prevent backflow into the thoracic cavity. The animal was then maintained undisturbed for approximately 90 minutes to allow the compound to harden.

After perfusion with Microfil or Vascupaint, mice were scanned using a Skyscan 1276 micro-CT system (Bruker) or a U-CT optical imaging system (MILabs). Images were acquired at a pixel size of 20 μm and a resolution of 2,016 × 1,344 pixels. Micro-CT scans were reconstructed using NRecon software (Bruker), with corrections for beam-hardening and ring artifacts. Image sets were saved as DICOM or BMP files (∼1,200–1,500 images per animal). Three-dimensional reconstructions were generated using 3D Slicer, and all non-aortic regions were removed using the scissors tool.

Aortic width and volume were quantified using the Segment Statistics module in 3D Slicer. For regional analyses, the aorta was divided into four segments: (1) ascending aorta and arch, extending from the sinotubular junction to immediately distal to the left subclavian artery; (2) descending thoracic aorta, extending from immediately distal to the left subclavian artery to the 13th rib pair; (3) suprarenal and renal aorta, extending from below the 13th rib pair to immediately distal to the left renal artery; and (4) infrarenal aorta, extending from immediately distal to the left renal artery to 1 mm into each iliac branch.

### Ultrasonography

Luminal diameters of the infrarenal aorta were measured using a Vevo 3100 ultrasound system equipped with an MS550 transducer (FUJIFILM VisualSonics Inc.). Mice were anesthetized with isoflurane (1-2% vol/vol) to maintain heart rates between 400 and 550 beats/minute and placed on a heated platform (37 °C) to prevent hypothermia. Color Doppler imaging was used to confirm pulsatile blood flow, and the probe was moved caudally to identify the right and left renal arteries. A cine loop of the infrarenal aorta distal to the left renal artery was acquired in the transverse view to determine the maximal luminal diameter. Maximal luminal diameters were measured from the acquired images using Vevo LAB 3.1.1 software (FUJIFILM VisualSonics Inc.).

### Western Blotting

After removal of the endothelium and adventitia, aortic tissues were homogenized in T-PER buffer (Catalog #: 78510; Thermo Scientific) supplemented with a protease inhibitor cocktail (Catalog #: P8340; Sigma-Aldrich) using disposable pellet pestles (Catalog #: Z359971; DWK Life Sciences LLC). Protein concentrations were determined using a BCA assay kit (Catalog #: 23225; Thermo Scientific). Protein samples were separated by SDS-PAGE and electrophoretically transferred to PVDF membranes. After blocking, the membranes were incubated with primary antibodies against LRP1 and β-actin, followed by appropriate secondary antibodies. Antibody information is provided in the Major Resources Tables. LRP1 and β-actin were detected by chemiluminescence (Catalog #: 34080; Thermo Fisher Scientific) using a ChemiDoc imaging system (Catalog #: 12003154; Bio-Rad).

### Histology

Infrarenal aortas were harvested, fixed in 10% (wt/vol) neutral-buffered formalin, and subsequently incubated in 70% (vol/vol) ethanol. Aortic tissues were paraffin-embedded and sectioned at 5 µm using a microtome. Paraffin-embedded sections were deparaffinized with limonene (Catalog #: 183164; Millipore-Sigma). Verhoeff iron hematoxylin staining was used to visualize elastic fibers.

Immunostaining for LRP1 and α-smooth muscle actin (α-SMA) was performed using the primary and secondary antibodies listed in the Major Resources Tables. Three negative controls were included: (1) omission of both primary and secondary antibodies, (2) secondary antibody only, and (3) substitution of the primary antibody with nonimmune rabbit IgG. NovaRed (Catalog #: SK-4805; Vector) was used as the chromogen. Histological and immunostaining images were acquired using an Axioscan 7 scanner (Zeiss) and analyzed with ZEN 3.1 Blue Edition software (Zeiss).

### Statistical Analysis

Data are presented as medians with 25^th^and 75^th^percentiles or as means, as appropriate for the statistical analysis performed. Two-group comparisons were conducted using the Mann–Whitney U test in SigmaPlot version 16 (SYSTAT Software Inc.). Log-transformed ultrasound data were analyzed using a mixed-effects model with random intercepts and random slopes for time (week), implemented with the nlme package in R version 4.4.2. Statistical significance was defined as P<0.05.

## RESULTS

### LRP1 Deletion in SMCs Promoted Extensive Aortic Dilatations, with Predominant Aneurysm Formation in the Infrarenal Abdominal Aorta

We first confirmed that *Sm22-*Cre-mediated deletion of LRP1 in SMCs did not affect other cell types within the aortic wall. Western blotting demonstrated a marked reduction in LRP1 protein abundance in aortic tissues from SMC-LRP1-/-mice after removal of the endothelium and adventitia (**Figure S1B**). Immunostaining of aortic tissue sections confirmed loss of LRP1 in the medial layers, whereas LRP1 in the adventitia was preserved in SMC-LRP1-/-mice (**Figure S1C**). Thus, consistent with previous reports,^4^LRP1 deletion was restricted to SMCs and did not involve other cell types within the aortic wall.

We next used micro-CT to visualize the entire aorta in 40-week-old mice. In both sexes, SMC-LRP1-/-mice exhibited increased aortic width across all four regions (**Figures 1B, S2, and S3; Table S1**), although the increase in the ascending aorta of male mice did not reach statistical significance. The greatest dilatation relative to SMC-LRP1+/+ littermates occurred in the infrarenal aorta, which was enlarged approximately 4-fold in males and 7-fold in females (**Figures 1B and S3**).

**Figure 1.**
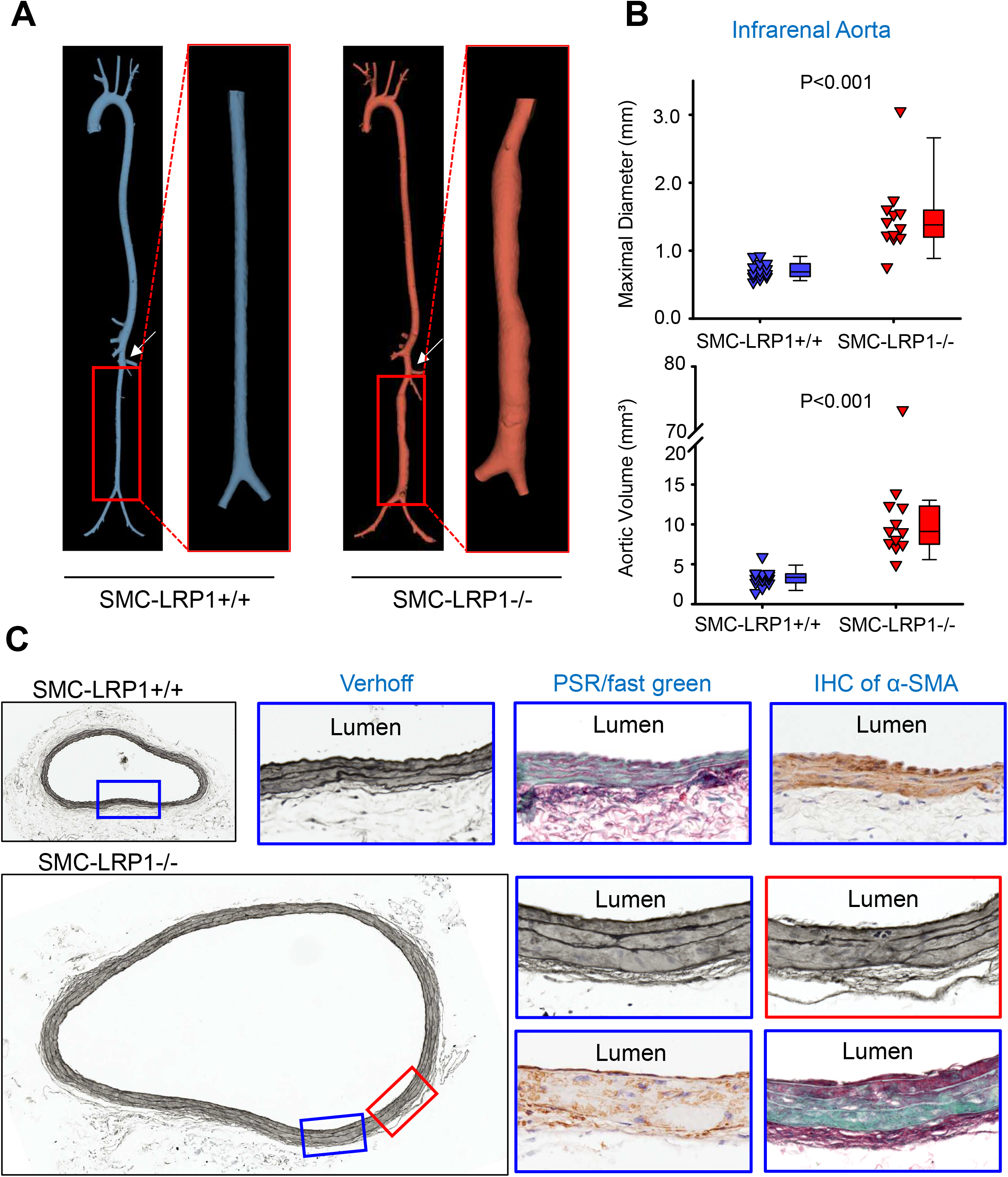
Deletion of LRP1 in SMCs induced infrarenal aortic aneurysms. Three-dimensional (3D) reconstructions of micro-CT images of the entire aorta from male 40-week-old SMC-LRP1+/+ and SMC-LRP1-/-mice. **(A)** Representative 3D images of the entire aorta. White arrows indicate the left renal artery. **(B)** Maximal diameter and volume of the infrarenal aorta in male mice. N = 12-14/group. Statistical analysis was performed using the Mann–Whitney rank-sum test. **(C)** Verhoeff staining, picrosirius red/fast green staining, and α-SMA immunostaining of infrarenal aortas.

At termination, histological analysis revealed profound SMC loss, as indicated by reduced α-SM actin immunostaining, together with marked collagen deposition in the medial layers and elastic fiber fragmentation (**Figure 1C**).

### Inhibition of Hepatocyte-Derived AGT Attenuated Infrarenal Aortic Dilatation

Because aortic dilatation was most pronounced in the infrarenal region, subsequent analyses focused on this segment. To determine when pathological changes emerged, we examined the infrarenal aorta histologically in 4-week-old mice (**Figure 2 and Figure S4**). At this age, neither overt SMC loss nor collagen deposition was evident; however, elastic fiber fragmentation was already present (**Figure 2A**).

**Figure 2.**
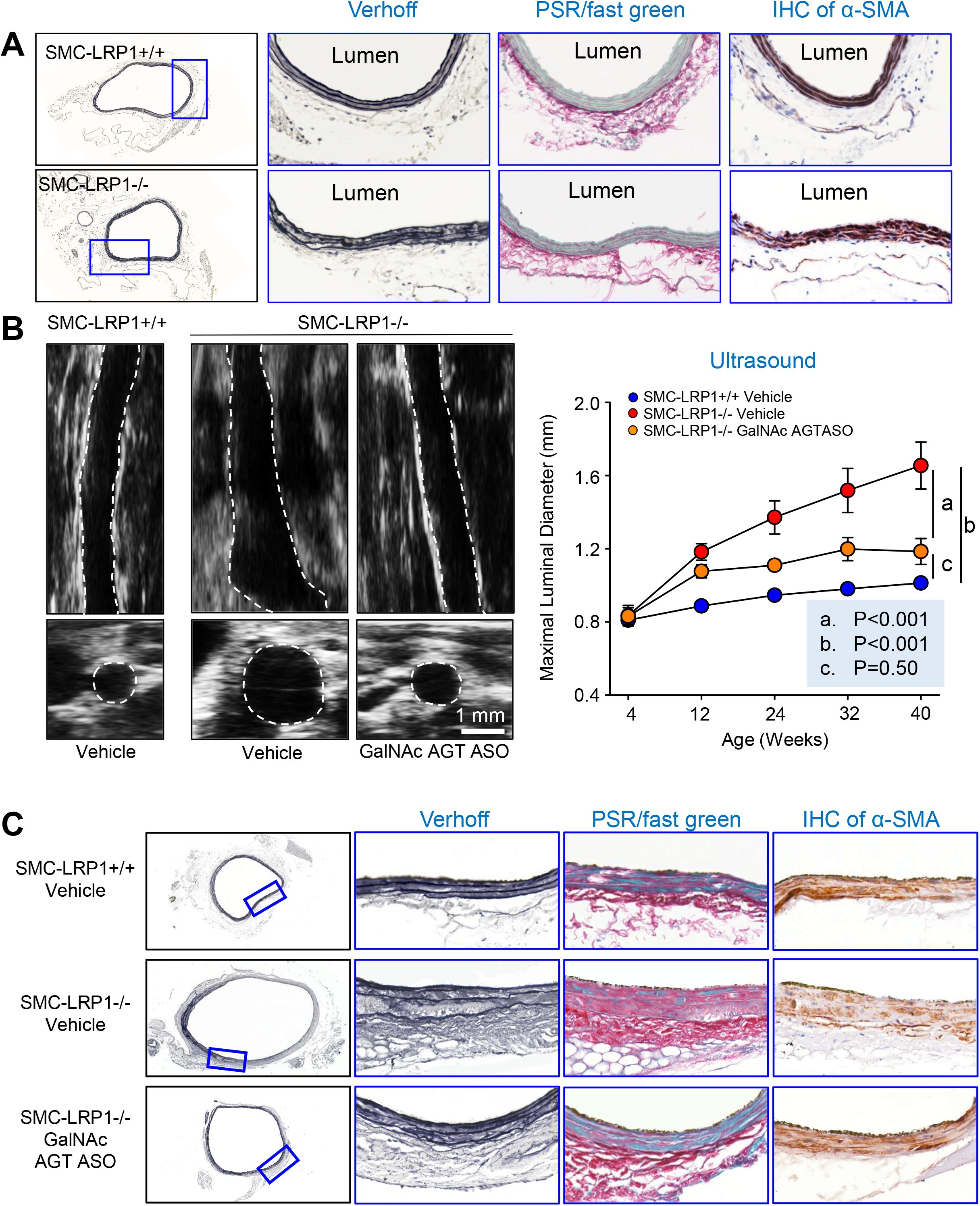
Loss of LRP1 in SMCs caused progressive infrarenal aortic luminal dilatation in male mice, which was attenuated by GalNAc AGT ASO. **(A)** Verhoeff staining, picrosirius red (PSR)/fast green staining, and α-SMA immunostaining of infrarenal aortas from 4-week-old male mice. **(B)** Ultrasonographic measurements of infrarenal aortic luminal diameters from 4 to 40 weeks of age in SMC-LRP1+/+ and SMC-LRP1-/-male mice administered vehicle or GalNAc AGT ASO. N = 9-10/group. Statistical analysis was performed using a mixed-effects model with random intercepts and random slopes for time (week). **(C)** Verhoeff staining, PSR/fast green staining, and α-SMA immunostaining of infrarenal aortas from 40-week-old male SMC-LRP1+/+ and SMC-LRP1-/-mice administered vehicle or GalNAc AGT ASO.

Despite modest disruption of the aortic wall, ultrasonography confirmed no apparent luminal dilatation at 4 weeks of age. This early stage, before evident luminal expansion, provided an opportunity to determine whether inhibiting AGT, the sole precursor of renin–angiotensin peptides, could prevent infrarenal aortic dilatation. Because our previous studies established that AGT is derived predominantly from hepatocytes,^19^4-week-old male SMC-LRP1-/-mice received weekly injections of vehicle or a GalNAc AGT ASO targeting hepatocyte-derived AGT until 40 weeks of age. Vehicle-injected male SMC-LRP1+/+ littermates served as normal controls. Ultrasonography was used to monitor luminal diameters at baseline, before injections began, and at 12, 24, 32, and 40 weeks of age (**Figure 2B and Figure S4**). Baseline luminal diameters did not differ among the three groups. In vehicle-injected SMC-LRP1-/-mice, infrarenal aortic luminal diameters increased progressively over time. Although luminal diameters also increased from baseline in GalNAc AGT ASO-administered SMC-LRP1-/-mice, inhibition of liver-derived AGT reduced the progression of luminal dilatation.

Luminal diameters did not differ significantly between GalNAc AGT ASO-injected SMC-LRP1-/-mice and vehicle-injected SMC-LRP1+/+ littermates. Histological analysis demonstrated greater SMC preservation and less collagen deposition in GalNAc AGT ASO-treated SMC-LRP1-/-mice than in vehicle-injected SMC-LRP1-/-mice (**Figure 2C**), indicating improved aortic wall integrity.

## DISCUSSION

Previous studies of SMC-LRP1-/-mice reported progressive aortic root dilatation^4^and superior mesenteric artery remodeling;^8^therefore, our initial objective was to determine whether loss of SMC-derived LRP1 causes additional aortic pathologies. Consistent with prior observations, SMC-LRP1-/-mice exhibited diffuse aortic dilatations. More importantly, micro-CT revealed marked, progressive dilatation of the infrarenal aorta in both sexes. This finding is clinically relevant because the infrarenal aorta is the predominant site of human AAAs and genetic variants in *LRP1* are associated with AAA risk. To our knowledge, the only other mouse strains reported to develop spontaneous or angiotensin II (AngII)-induced infrarenal AAAs are those with inducible deletion of YAP/TAZ in SMCs^20^, liver kinase B1 in SMCs,^21^or inducible CCN2 in SMCs,^15^although their pathological features remain incompletely characterized.

The infrarenal localization of aneurysm formation in SMC-LRP1-/-mice is particularly notable, as most commonly used mouse models with genetic and pharmacological manipulations do not have overt dilatation in this regional distribution. AngII infusion induces AAAs primarily in the suprarenal abdominal aorta, without reported pathological involvement of the infrarenal region.^14, 22, 23^Similarly, systemic aldosterone administration,^24^as well as co-administration of AngII with BAPN^25^or a TGFβ-neutralizing antibody,^26^induces AAAs that are confined to the suprarenal aorta, with no evident infrarenal involvement. In our previous study, AngII infusion augmented dilatation in the ascending aorta, but not the suprarenal abdominal aorta, in 8- to 12-week-old male SMC-LRP1-/-mice; however, infrarenal aortic diameters were not measured.^5^The current findings therefore extend these prior observations by demonstrating that SMC-LRP1 deficiency promotes progressive luminal dilatations in the infrarenal aorta. The basis for the apparent regional resistance or susceptibility of specific aortic segments in different mouse studies described above remains unclear. Interestingly, in contrast to AngII-induced AAA development, intraperitoneal injection of Lactobacillus casei cell-wall extract in a mouse model of Kawasaki disease induces aneurysms that localize primarily to the infrarenal aorta.^27^Together, these observations highlight the complexity of region-specific aortic pathobiology and imply that distinct mechanisms may selectively target the infrarenal aorta under specific conditions.

Sex differences are an important consideration in AAA biology. In humans, AAAs occur more frequently in men than in women, and a similar male predominance is observed in AngII-induced AAAs in mice.^28, 29^In the present study, both male and female SMC-LRP1-/-mice developed infrarenal AAAs, indicating that SMC-LRP1 deficiency promotes AAAs in both sexes. However, because male and female mice were not directly compared side by side within the same experimental cohort, whether sex influences the severity, progression, or underlying mechanisms of infrarenal AAAs caused by SMC-LRP1 deficiency remains to be determined. Adequately powered sex-specific comparisons will be important for defining whether loss of SMC-LRP1 interacts with sex-dependent pathways to promote AAA development and progression.

After defining the infrarenal phenotype in SMC-LRP1-/-mice, we next explored whether the renin-angiotensin system contributes to aneurysm progression. Although AngII infusion does not induce infrarenal AAAs in wild-type mice,^14, 22, 28^recent studies showed that AngII promotes infrarenal AAAs in mice with inducible CCN2 deletion in SMCs, with no or only modest pathology in the suprarenal aorta.^15^This observation is relevant because CCN2 is a known ligand of LRP1.^7^Therefore, disruption of LRP1-related signaling in SMCs may create a permissive environment in which AngII contributes to infrarenal AAA development and progression. Because AGT is the sole substrate for AngII generation and is produced primarily by hepatocytes, we determined whether pharmacological inhibition of liver-derived AGT would attenuate infrarenal AAAs in SMC-LRP1-/-mice. This approach also has translational relevance, as inhibition of hepatocyte-derived AGT with ASO or small interfering RNA has been evaluated in clinical trials.^30-32^Our data show that GalNAc AGT ASO slowed progressive infrarenal aortic dilatation and improved aortic wall integrity, supporting a contribution of hepatocyte-derived AGT and AngII signaling to aneurysm progression caused by SMC-LRP1 deficiency.

In summary, deletion of LRP1 in SMCs causes extensive aortic dilatation, with spontaneous aneurysm formation most prominently affecting the infrarenal aorta. This phenotype provides a clinically relevant research tool for studying infrarenal AAA pathobiology and molecular mechanisms. Pharmacological inhibition of hepatocyte-derived AGT attenuated aneurysm progression, suggesting that AGT-dependent AngII signaling contributes to disease progression in the setting of SMC-LRP1 deficiency. Future studies will focus on defining the molecular mechanisms linking loss of SMC-derived LRP1 to infrarenal aneurysm formation and progression.

## Non-standard Abbreviations

AGT: Angiotensinogen
AngII: Angiotensin II
ASO: Antisense oligonucleotides
GalNAc: N-acetylgalactosamine
LRP1: Low-density lipoprotein receptor-related protein 1
Micro-CT: Micro-computed tomography
SMC: Smooth muscle cells

## ACKNOWLEDGMENTS

Micro-CT was performed using SkyScan 1276 micro-CT at the Magnetic Resonance Imaging and Spectroscopy Center (RRID:SCR_026383) or a U-CT Optical Imaging system (MILabs) at the Light Microscopy Core (RRID:SCR_026405) at the University of Kentucky. Histological and immunostaining images were taken using Zeiss Axioscan 7 in the Light Microscopy Core at the University of Kentucky (RRID:SCR_026405).

## SOURCES OF FUNDING

The authors’ research work was supported by the National Heart, Lung, and Blood Institute of the National Institutes of Health (R35HL155649), the American Heart Association MERIT award (23MERIT1036341), and the Leducq Foundation for the Networks of Excellence Program (Cellular and Molecular Drivers of Acute Aortic Dissections).

## DISCLOSURES

Alan Daugherty and Hong S. Lu hold a patent for the use of antisense oligonucleotides targeting angiotensinogen in thoracic aortic aneurysms in patients with Marfan syndrome. Adam E. Mullick is an employee of Ionis Pharmaceuticals, which provided the mouse GalNAc AGT ASO. The other authors declare that they have no conflicts of interest.

## SUPPLEMENTAL MATERIALS

Supplemental Figures 1-4 Major Resources Tables

