## Supplemental Materials for "Deletion of LRP1 in Smooth Muscle Cells Induces Infrarenal Aortic Aneurysms in Mice"

Michael K. Franklin,^1*^ Guan Fan,^1*^ Bowen Li,^1*^ Pengjun Wang,^1^ Deborah A. Howatt,^1^

Jessica J. Moorleghen^1^, Adam E. Mullick,^3^ Alan Daugherty,^1,2^ Hong S. Lu^1,2^

^1^ Saha Cardiovascular Research Center and Saha Aortic Center

^2^ Department of Physiology

University of Kentucky, Lexington, Kentucky, USA

^3^ Ionis Pharmaceuticals, Inc., Carlsbad, CA, USA

*These authors contributed equally

**Running Title:** LRP1 and Infrarenal Aortic Aneurysm

**Corresponding Authors:**

Alan Daugherty

Hong S. Lu

**MAJOR RESOURCES TABLES**

**Mouse Models**

| **Strain** | **Vendor or Source** | **Background Strain** | **Sex** | **Other Information (e.g. breeding scheme for F1, F2…if applicable)** | **Persistent ID / URL** |
| --- | --- | --- | --- | --- | --- |
| *Lrp1^fl/fl^* | JAX (Strain #012604) | N/A | F | This strain was purchased for mouse breeding to generate cell-specific deletion of *Lrp1* | https://www.jax.org/strain/012604 |
| *Sm22-Cre^+/0^* | JAX (Strain# 017491) | C57BL/6J | M | This strain was purchased for mouse breeding to generate *Lrp1* deletion in smooth muscle cells (SMCs) | https://www.jax.org/strain/017491 |
| *Lrp1^fl/f^Sm22-Cre* | In house | C57BL/6J  (N: N/A) | M&F | Breeding Scheme:  **F0:** Female *Lrp1^fl/fl^* x Male *Sm22-Cre^+/0^*  **F1:** Female *Lrp1^fl/+^Sm22-Cre^0/0^* x Male *Lrp1^fl/+^ Sm22-Cre^+/0^*  **F2:** Female *Lrp1^fl/fl^ Sm22-Cre^0/0^* x Male *Lrp1^fl/fl^ Sm22-Cre^+/0^* |  |

**Primary Antibodies for Immunostaining**

| **Antibody** | **Vendor** | **Cat #** | **Working Concentration** |
| --- | --- | --- | --- |
| Rabbit anti-LRP1 | abcam | Ab92544 | 1 µg/mL |
| Rabbit nonimmune IgG | ImmunoReagents | Rb-003-V | 1 µg/mL |

**Secondary Antibodies for Immunostaining**

| **Antibody** | **Vendor** | **Cat #** | **Working Concentration** |
| --- | --- | --- | --- |
| ImmPress Goat Anti-Rabbit IgG | Vector | MP-7451 | No concentration information available; Ready-to-use (per manufacturer) |

**Animal Study Information Following the ARRIVE Essential 10**

**Figure 1 and Figure S3**

| **Groups** | **Sex** | **Age (weeks)*** | **Number**  **(Start)** | **Number (Termination)** |
| --- | --- | --- | --- | --- |
| SMC-LRP1+/+ | Male | 40 | 17 | 15 |
| SMC-LRP1-/- | Male | 40 | 27 | 24 |
| SMC-LRP1+/+ | Female | 40 | 17 | 16 |
| SMC-LRP1-/- | Female | 40 | 32 | 32 |

**Figure 2C**

| **Groups** | **Sex** | **Age (weeks)*** | **Number**  **(Start)** | **Number (Termination)** |
| --- | --- | --- | --- | --- |
| SMC-LRP1+/+ | Male | 4 – 40 | 9 | 9 |
| SMC-LRP1-/- | Male | 4 – 40 | 9 | 9 |
| SMC-LRP1+/+ | Female | 4 – 40 | 6 | 6 |
| SMC-LRP1-/- | Female | 4 – 40 | 7 | 7 |

**Figure S1B**

| **Groups** | **Sex** | **Age (weeks)*** | **Number**  **(Start)** | **Number (Termination)** |
| --- | --- | --- | --- | --- |
| SMC-LRP1+/+ | Male | 4 | 4 | 4 |
| SMC-LRP1-/- | Male | 4 | 4 | 4 |

**Figure S2 Male**

| **Groups** | **Sex** | **Age (weeks)*** | **Number**  **(Start)** | **Number (Termination)** |
| --- | --- | --- | --- | --- |
| SMC-LRP1+/+ | Male | 40 | 12 | 10 |
| SMC-LRP1-/- | Male | 40 | 22 | 19 |

**Figure S2 Female and Figure S4**

| **Groups** | **Sex** | **Age (weeks)*** | **Number**  **(Start)** | **Number (Termination)** |
| --- | --- | --- | --- | --- |
| SMC-LRP1+/+ | Female | 40 | 13 | 12 |
| SMC-LRP1-/- | Female | 40 | 27 | 27 |

*Age (weeks): the age when the experiment was started (except Figure 2A) or the age range (Figure 2A).

**ARRIVE Essential 10 Checklist**

| **Item** | **Application** |
| --- | --- |
| Ethics | Approved by the University of Kentucky IACUC (2018-2967). |
| Sex | Described in each figure. |
| Inclusion criteria | Based on sex, age, body weight, and overall health appearance in each experiment. |
| Exclusion criteria | Based on medical cases reported by a veterinarian. |
| Sample size | Described in the above tables. |
| Sample size calculation | None |
| Primary endpoint | Aortopathies |
| Randomization | Study mice within each genotype were numbered and grouped randomly |
| Blinding | Quantification of aortic width, length, and volume (micro-CT) and luminal diameters (ultrasound) was measured by an investigator blinded to the study group information. Some data were measured by two investigators independently to validate the consistency of measurements. |
| Statistical analysis | SigmaPlot 16 (SYSTAT Software Inc., CA) or R Statistical Software. |
| Statistical method | Described in the Statistical Analysis Section. |
| Data availability | All numerical data used for figures are available in the Supplemental Excel File. |
