## Supplemental Figures for "Deletion of LRP1 in Smooth Muscle Cells Induces Infrarenal Aortic Aneurysms in Mice"

**A**

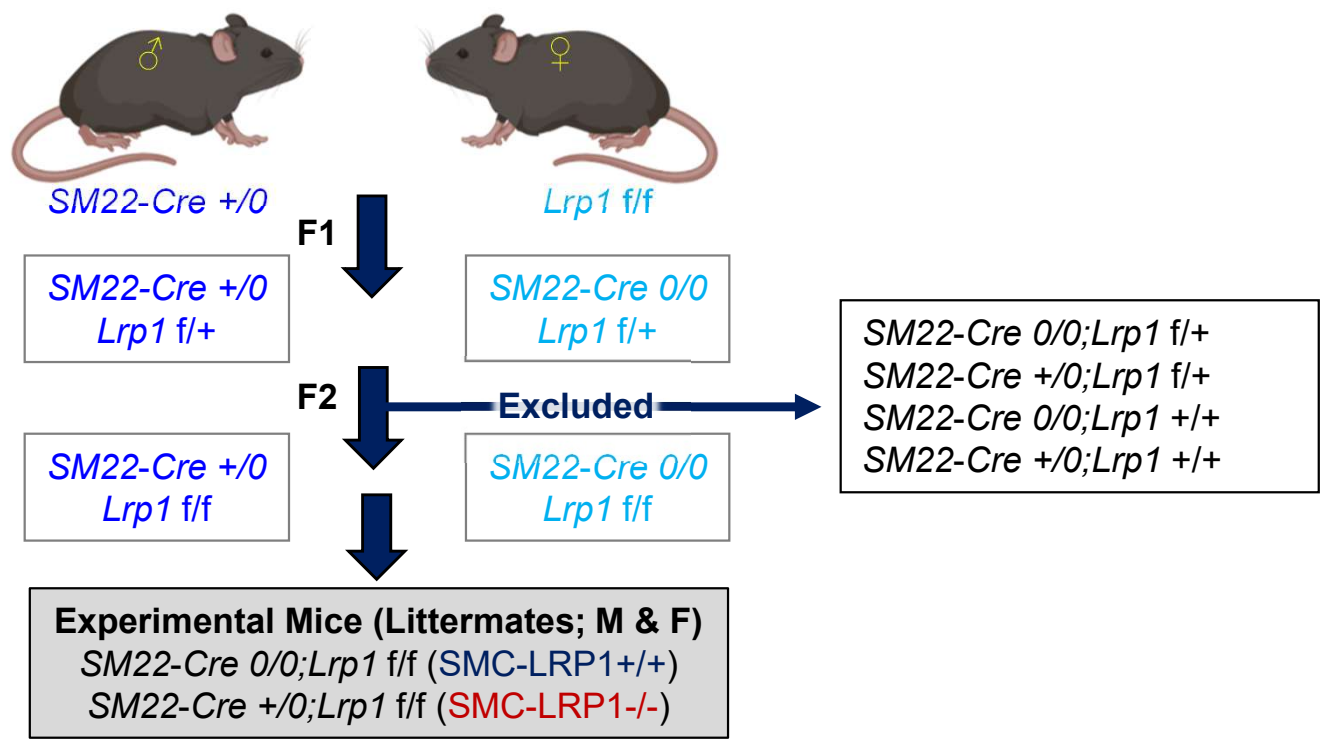

**B**

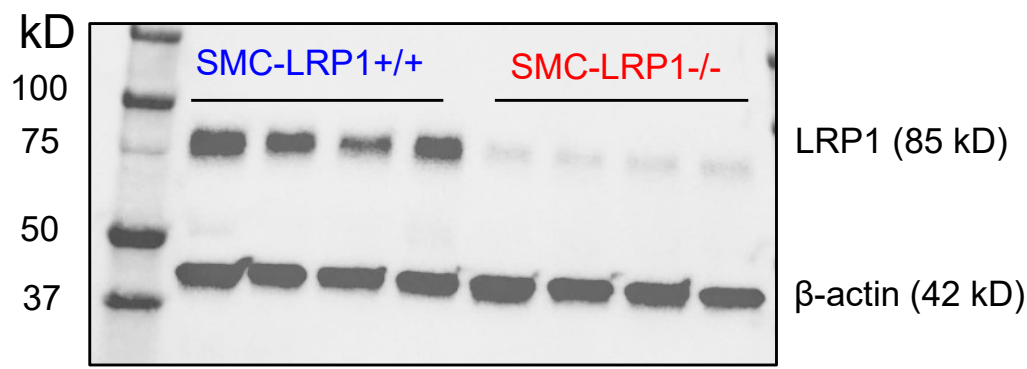

**C**

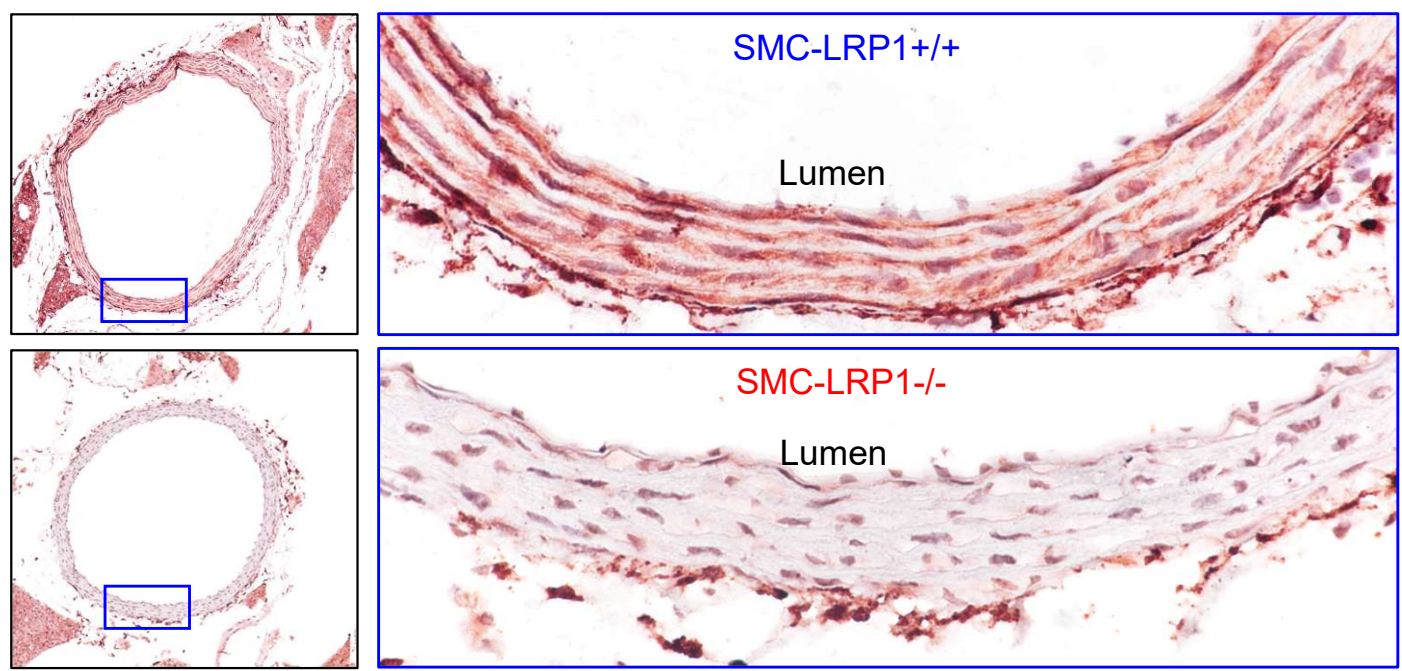

**Figure S1. Breeding strategy and confirmation of LRP1 deletion in smooth muscle cells (SMCs).** (A) Breeding strategy to develop SMC-LRP1<sup>+/+</sup> and SMC-LRP1<sup>-/-</sup> littermates. LRP1 deletion in aortic SMCs was confirmed by Western blot (B) and immunostaining of aortas (C) in SMC-LRP1<sup>-/-</sup> mice. Graphic mouse images were created with BioRender.

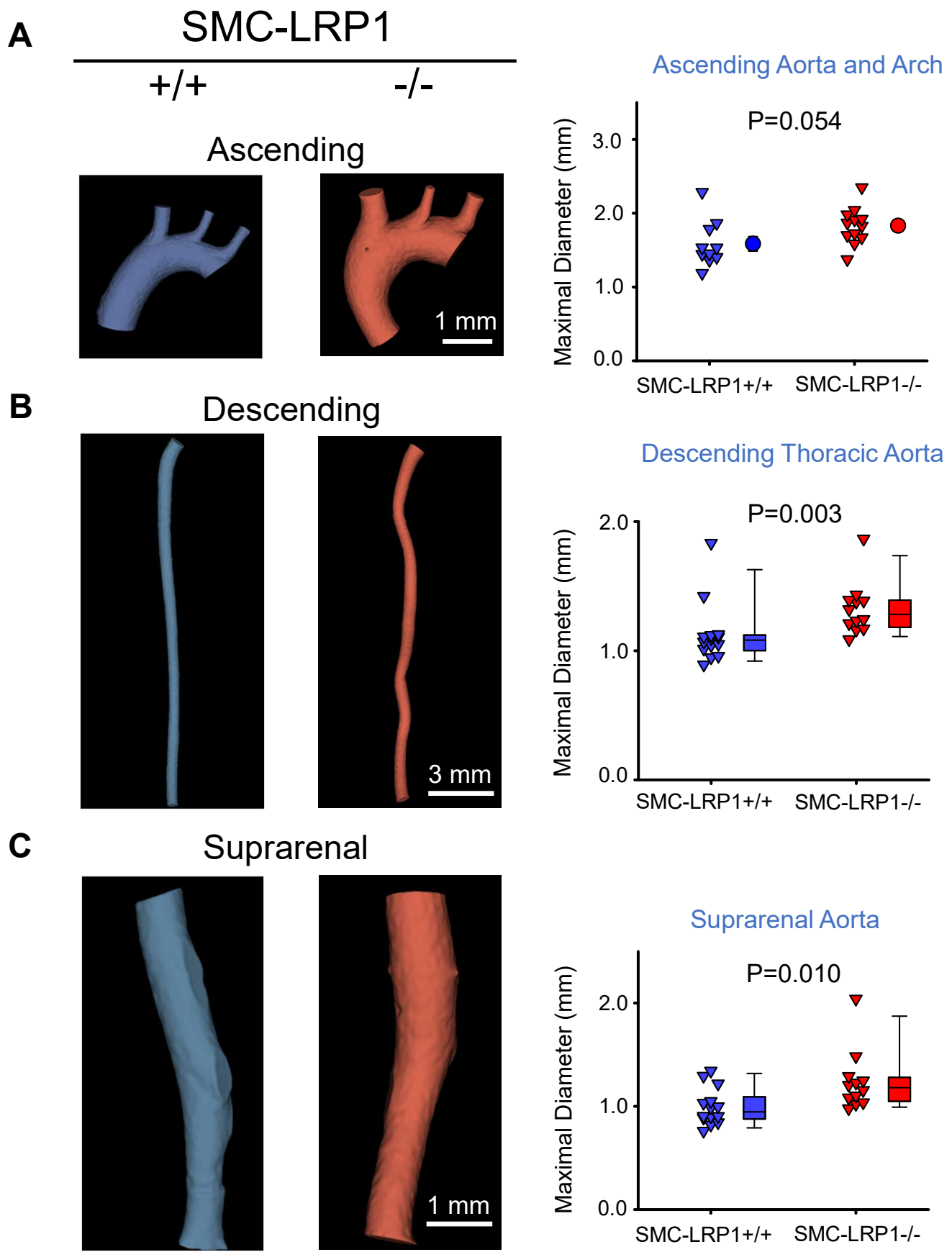

**Figure S2. LRP1 deletion in SMCs induced multiple aortic pathologies spontaneously in male mice.** 3D reconstruction of micro-CT images and diameter quantification of the ascending (A), descending thoracic (B), and suprarenal (C) regions in male SMC-LRP1<sup>+/+</sup> and SMC-LRP1<sup>-/-</sup> mice at 40 weeks of age. N =10 to 14/group. Statistical analysis: Student's t test (A) or Mann-Whitney rank sum test (B-C).

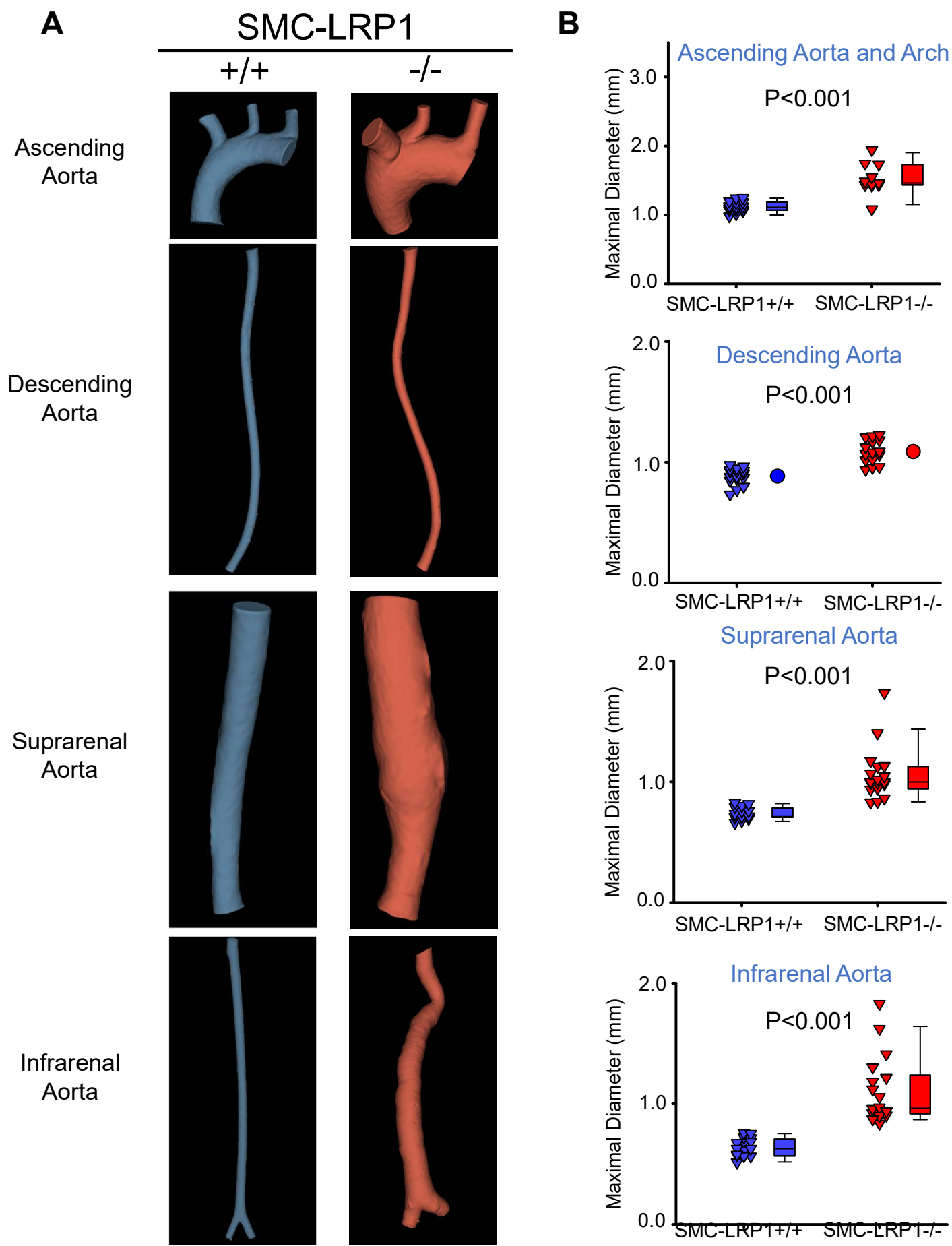

**Figure S3. LRP1 deletion in SMCs induced multiple aortic pathologies spontaneously in female mice.** 3D reconstruction of micro-CT images (**A**), and diameter quantification (**B**) of the ascending, descending, suprarenal, and infrarenal regions in female SMC-LRP1+/+ and SMC-LRP1-/- mice at 40 weeks of age. N=14 to 16/group. Student's t test (descending thoracic aorta) or Mann-Whitney rank sum test (other aortic regions).

**A**

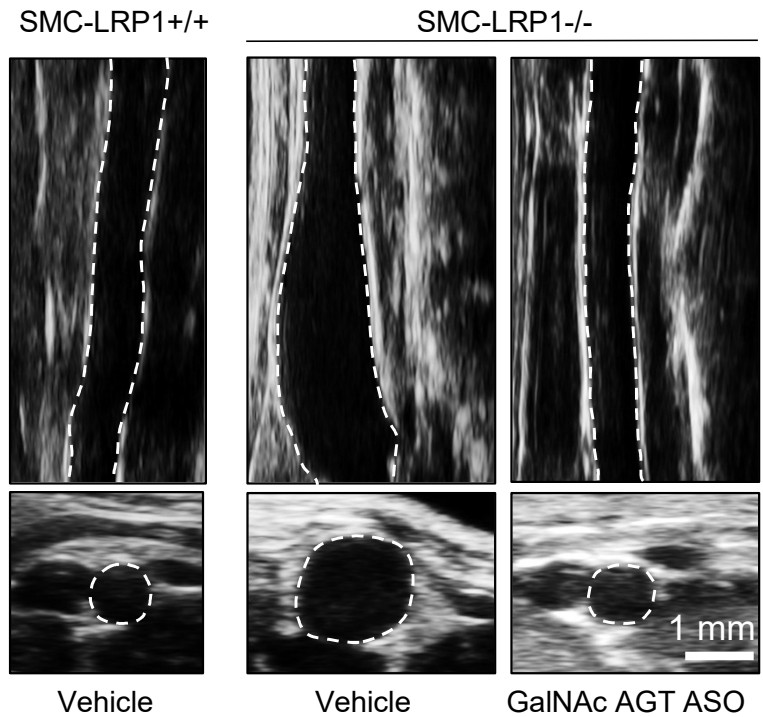

**B**

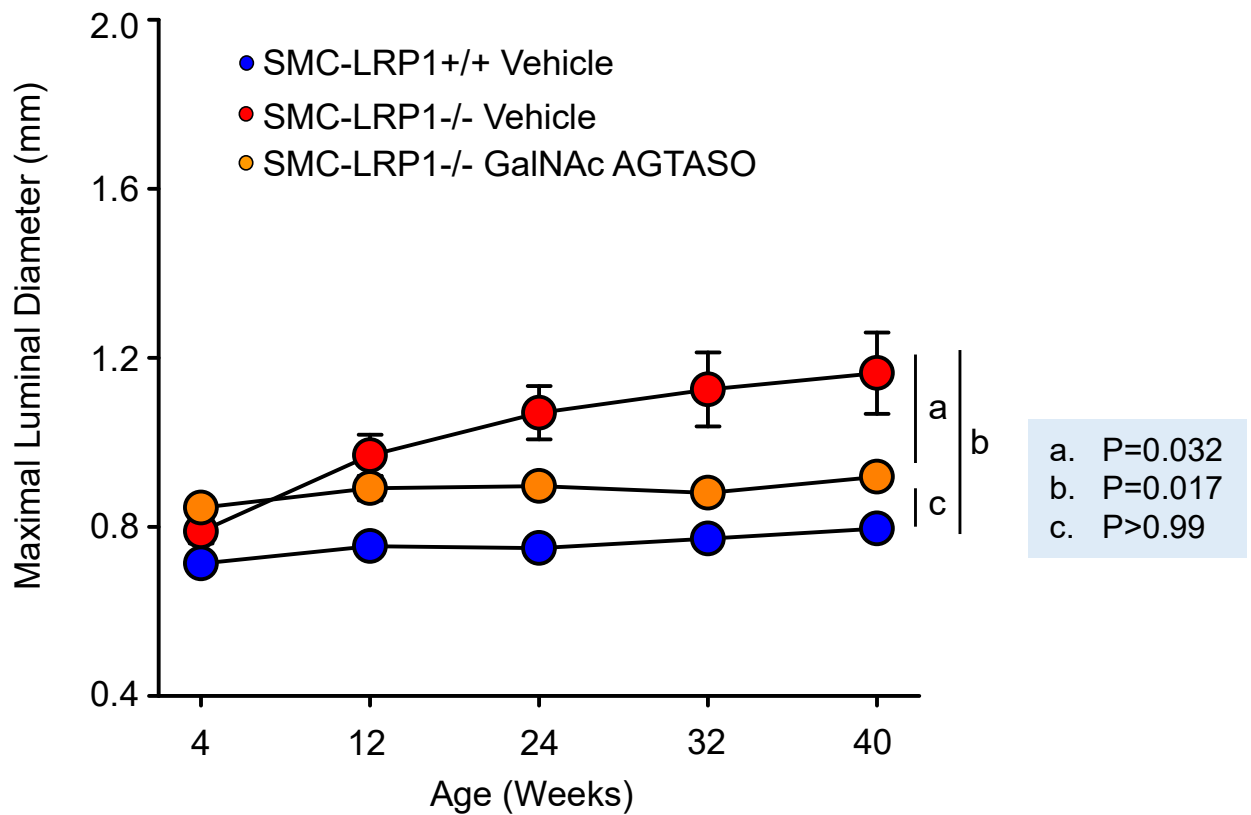

**Figure S4. Progressive infrarenal aortic luminal dilatations in female SMC-LRP1<sup>-/-</sup> mice were attenuated by GalNAc AGT ASO.** Representative ultrasound images of infrarenal aortas from 40-week-old SMC-LRP1<sup>+/+</sup> and SMC-LRP1<sup>-/-</sup> mice (**A**). Ultrasonographic measurements of infrarenal aortic luminal diameters from 4 to 40 weeks of age in SMC-LRP1<sup>+/+</sup> and SMC-LRP1<sup>-/-</sup> female mice administered vehicle or GalNAc AGT ASO. N = 9-10/group (**B**). Statistical analysis was performed using a mixed-effects model with random intercepts and random slopes for time (week).
